# Towards substitution of invasive telemetry: An integrated home cage concept for unobstrusive monitoring of objective physiological parameters in rodents

**DOI:** 10.1101/2023.05.12.540546

**Authors:** Lucas Mösch, Janosch Kunczik, Lukas Breuer, Dorit Merhof, Peter Gass, Heidrun Potschka, Dietmar Zechner, Brigitte Vollmar, René Tolba, Christine Häager, André Bleich, Michael Czaplik, Carina Barbosa Pereira

**Affiliations:** Department of Anaesthesiology, Faculty of Medicine, RWTH Aachen University, Aachen, North Rhine-Westphalia, Germany; Chair of Image Processing, Faculty of Computer and Data Science, Universität Regensburg, Regensburg, Bavaria, Germany; RG Animal Models in Psychiatry, Department of Psychiatry and Psychotherapy, Central Institute of Mental Health Mannheim, Medical Faculty Mannheim, University of Heidelberg, Heidelberg, Baden Württemberg, Germany; Institute of Pharmacology, Toxicology, and Pharmacy, Ludwig-Maximilians-University, Munich, Bavaria, Germany; Rudolf-Zenker-Institute of Experimental Surgery, University Medical Centre Rostock, Rostock, Mecklenburg-Western Pomerania, Germany; Institute of Laboratory Animal Science, Faculty of Medicine, RWTH Aachen University, Aachen, North Rhine-Westphalia, Germany; Institute for Laboratory Animal Science, Hannover Medical School, Hannover, Lower Saxony, Germany

**Author notes:** These authors contributed equally to this work (Lucas Mösch, Janosch Kunczik).

## Abstract

This study presents a novel concept for a smart cage designed to monitor the physiological parameters of mice and rats in animal-based experiments. The system focuses on monitoring key clinical parameters, including heart rate, respiratory rate, body temperature, activity, and circadian rhythm. To create the smart home cage system, an in-depth analysis of the requirements was performed, including camera positioning, imaging system types, resolution, frame rates, external illumination, video acquisition, data storage, and synchronization. Two different camera perspectives were considered, and specific camera models, including two near-infrared and two thermal cameras, were selected to meet the requirements. During the first testing phase, the system demonstrated the potential of extracting vital parameters such as respiratory and heart rate. This technology has the potential to reduce the need for implantable sensors while providing reliable and accurate physiological data, leading to refinement and improvement in laboratory animal care.

## Introduction

The fundamental aim of biomedical research is to understand the etiology and pathophysiology of diseases so that strategies and recommendations might be developed for their prevention and treatment [1–3]. Animal experiments played a significant role in this understanding and contributed to the tremendous medical progress achieved to date [4]. However, ethical concerns regarding animal pain, distress, and death during scientific experiments have long been a highly controversial topic [5]. Several alternative methods have been proposed to overcome the drawbacks inherent in animal studies and to avoid unethical procedures. The 3Rs principle by Russell and Burch (1959) – Replace, Reduce, and Refine – has been widely incorporated into animal research laws and regulations to maintain high animal welfare standards [6]. These principles promote alternative methods and reduce the number of animals used in scientific research while refining the trials to ensure the most efficient and least severe procedures [5]. In addition, several methods, such as in vitro testing, computer – in silico – modeling, and human-patient simulators, have been proposed to avoid the use of animals [7, 8]. These methods are indeed an essential and integral part of research. However, they do not necessarily lead to a complete replacement of animal trials, as only very few experiments can (so far) be entirely substituted by alternative methods. Instead, these approaches exist parallel or complementary to each other [9].

The accuracy and transferability of animal-based research are inherently linked to the health and well-being of the animals used. Animals in distress, experiencing pain, or under excessive stress may exhibit modified physiological and psychological responses, significantly impacting the quality of data obtained [10]. As a result, investigators are advised to recognize situations and procedures that may adversely affect the animals. Early identification of signs of suffering enables prompt intervention and refinement of humane endpoints. Furthermore, recognizing and evaluating techniques and methods, such as environmental enrichment, is crucial to enhance the animals’ welfare and promote positive outcomes.

Whether behavioral or physiological, welfare indicators should be objective, species-specific, easily recognized, and practical [11, 12]. Multiple indicators are recommended to avoid interpretation errors and provide a comprehensive understanding of an animal’s welfare. Importantly, clear concepts how to combine these indicators to objectively assess the severity of procedures have been developed [13]. Appropriate welfare indicators both psychological and physiological may be divided in six ‘high level’ categories:

- **Appearance**, including body, coat and skin conditions (e.g., unkempt coat, porphyrin staining);
- **Body functions**, such as changes in body weight and temperature as well as changes in heart rate (HR), respiratory rate (RR) and character, blood pressure and stress hormone levels;
- **Environment** within the cage (such as nest quality, consistency of faeces);
- **Behaviours**, including social interaction, posture, gait, lameness, and undesirable behaviour (e.g., excessive licking, scratching, apathy or withdrawal, increased aggression, stereotypic behaviour);
- **Procedure-specific indicators** (for instance, tumour size in cancer experiments);
- **Free observations** (subjective analysis of the researcher) [14, 15].

To date, the assessment of welfare indicators has been sporadic in both day-to-day work and during animal trials. In addition, numerical scoring systems used to evaluate animal health and welfare rely on objective measurements and personnel assessments [16]. Telemetric sensors can provide continuous vital signal monitoring in chronic experiments. However, their surgical implantation is stressful and can restrict animal movement. Furthermore, they have a short lifespan due to being powered by single-use batteries, which limits their use in animal studies [17, 18]. Therefore, one of the holy grails of animal-based research is achieving the continuous and non-intrusive monitoring of the affective state and physiological indicators throughout experiments [19].

Although continuous 24/7 monitoring may seem utopian, there have been significant progresses in remote monitoring in the last two decades, especially with the increasing advances in artificial intelligence. These methods permit to assess objective clinical signs such as body temperature [20, 21],HR [22–24] RR [22, 24, 25], food intake [26], behaviour including social interaction [27–29], movement [25, 30] and facial expressions [31, 32].

This paper provides a novel concept for a smart cage that can monitor clinical parameters such as HR, RR, body temperature, movement, and circadian rhythm in mice or rats. For this, the current state-of-the-art approaches for unobstrusive monitoring are presented. Based on a comprehensive requirements analysis for an automated camera-based homecage monitoring system, a reference concept is presented for a homecage system that monitors HR, RR, and body temperature, along with its specifications and software. Additionally, a feasibility study was conducted to investigate the practicality of this reference concept. Finally, the necessary hardware schematics and the software solution are also provided to replicate and utilize the proposed system.

### Approaches for unobstrusive monitoring

The following section provides an overview and brief description of the algorithms presented in previous publications that can be utilized for estimating objective parameters of rodents in the home cage through imaging techniques.

### Assessment of respiratory rate

Approaches for RR assessment of anesthetized rodents have been proposed by Pereira et al. (2018) [25] and Kunczik et al. (2019) [24] using infrared thermography (IRT) and visual imaging (VIS), respectively. These so-called motion-based approaches track the motion of explicit image regions over time and determine the trajectories that correlate with RR. These approaches consist of multiple steps:

- Finding the region of interest (thorax and abdomen) in the animal;
- Detecting and tracking of movement within the regions to measure the expansion and contraction occurring during the respiratory cycle;
- Extracting and filtering the resulting trajectories;
- Applying signal analysis to reduce noise and whiten the observations;
- Computing the frequency spectra, and thus the respiratory rate is derived from the frequency maximum;

However, it is strongly recommended to extract the respiratory rate only when the animal is still or asleep, as even slight disturbances, such as animal movements, can severely affect the spectra.

### Assessment of heart rate

The monitoring of heart activity can be achieved through VIS, IRT, and near-infrared (NIR) imaging. Each method requires a unique approach to estimating HR. VIS and NIR can be used for motion-based or intensity-based approaches to extract heart rates from videos, as the initial findings of Kunczik et al. (2019) [24] and Zhao et al. (2013) [33] suggest. Hereby, motion-based approaches detect body movements caused by the beating heart. The trajectories of these movements are processed similar to the RR assessment. The trajectories are tracked and filtered, and then signal analysis is used to reduce noise and whiten the data. HR is then determined by the highest frequency peak in the frequency spectrum.

In contrast, intensity-based approaches use changes in skin color or brightness due to blood flow to quantify heart activity. This method involves identifying and tracking ROI enclosing the animals’ skin and requires multiple steps:

- Finding the regions not covered by fur (ears, face, tail) in the animal;
- Detecting and tracking of color changes over time within these regions;
- Extracting and filtering the resulting signals;
- Applying signal analysis to reduce noise and whiten the observations;
- Computing the frequency spectra, and thus the RR is derived from the frequency maximum;

To ensure accurate measurement with these approaches, it is strongly recommended only to collect HR data from animals in their home cage while they are still or asleep. This is to avoid motion artifacts that could distort the entire measurement.

### Body and core temperature assessment

General temperature assessment can be performed easily using single frames of radiometric cameras. Through image object detection, it is possible to identify and track the animal’s body areas and calculate the average temperature of the skin area in the thermographic frames. Vinne et al. (2020) [21] proposed an approach to derive core temperature patterns from the temperature of the skin area assessed in this way. In addition, a method was proposed by van der Vogel et al. (2016) [34] for direct core temperature assessment. This approach uses the eye region to derive the core temperature from a thermographic image, requiring the eye to be prominently visible inside the captured image.

### Tracking of movement and circadian rhythm

In 2018, our research group presented an approach to evaluating both exploratory behavior and general activity in rodents using thermal imaging, which also be applied to visual and near-infrared imaging videos. A detailed description of the algorithm can be found in Pereira et al. (2018) [25]. During the rodents’ tracking process, their position and velocity are assessed continuously, resulting in cumulative time heat maps that reveal time spent by the animals in different parts of the cage. Analyzing the parameters of position, velocity, and time can provide insights into the circadian rhythm of the animals, including information about their sleep, drinking, daily activity, and more. In addition, this data can be used to compare and analyze differences in individual behavior patterns as well as between groups of animals and the effects of various experimental conditions or interventions.

### Technical requirement analysis

In order to enable the measurement of parameters through the described approaches, the home cage system must comply with specific technical requirements. Thus a comprehensive analysis of the requirements is needed for effective animal monitoring.

### Requirements for heart rate assessment

#### Camera positioning

The perspective used is crucial for assessing HR, as it determines which parts of the animals are best visible. The ideal camera position for intensity-based measurements in a rodent is frontal, providing an unobstructed view of the nose and ears. Although, it has limitations regarding occlusions and continuous assessments. For this purpose, a top-down birds-eye perspective is more appropriate as it provides an occlusion-free view of all animals. Therefore, both perspectives should be covered by a monitoring cage. Since the frontal camera can only cover a small portion of the cage, it is recommended to place it near a water or food source, where the animals spend a considerable amount of time. This way, the chance of obtaining frontal images of the rodents is maximized.

#### Camera sensor

For intensity-based HR assessment, hemoglobin is the primary source of pulse wave-induced changes. Maximum absorption and detectable intensity changes occur around 400 nm in the violet light spectrum, as shown in Fig 1 (left). While UV cameras are available that are sensitive to this spectral range (Fig 1 (right)), their high cost and intense UV light for optimal signal quality are not advisable due to increased radiation energy. Moreover, the monitoring system requires cameras to operate in low-light situations without ambient light. This requirement eliminates the possibility of utilizing green channels in conventional cameras. Therefore, NIR cameras, in combination with external NIR light, are best suited for monitoring systems. NIR light can penetrate deep into the tissue, enabling visualization of corresponding changes in hemoglobin concentration intensity. Additionally, research has indicated that rodents do not perceive NIR wavelengths between 700 and 900 nanometers and pose no risk to the animals.

**Fig 1.**
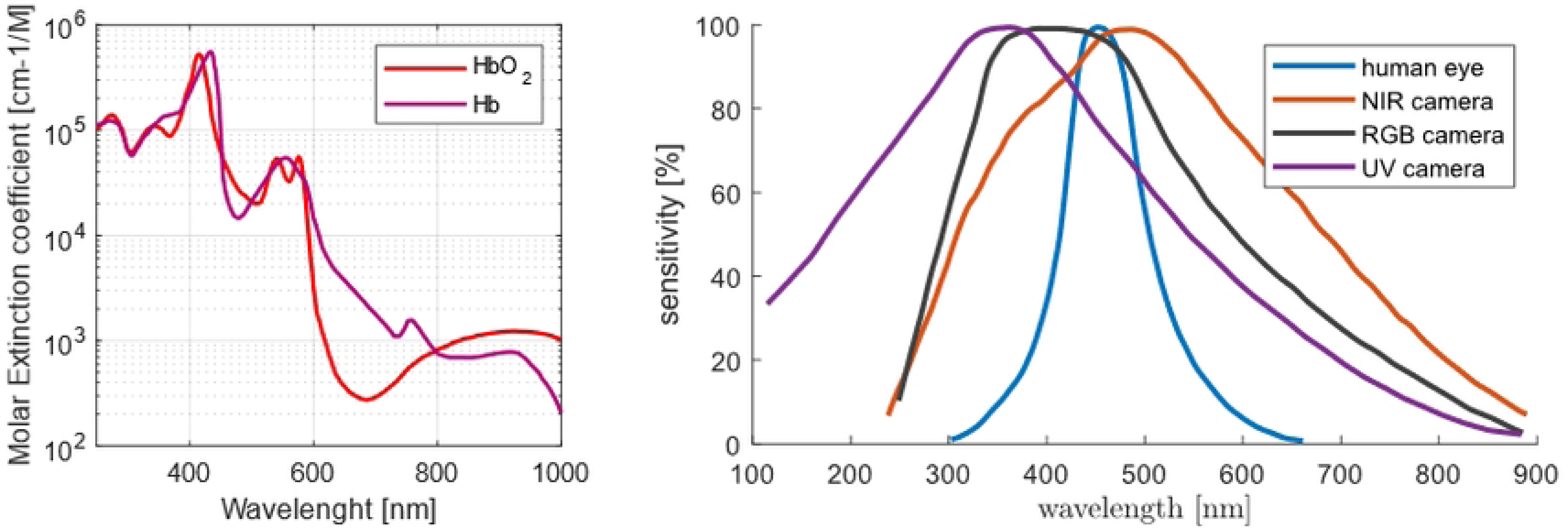
Light spectra relevant to intensity based HR assessment. Typical hemoglobin light absorption spectrum (left) compared to typical optical sensors sensitivity (right).

#### Frame rates

Frame rates are an important consideration in contactless HR measurement in rodents. The proper frame rate must ensure both the accuracy of the HR measurements and the feasibility of the system. According to the sampling theorem, to accurately measure a signal, the sampling rate must be at least twice the highest frequency component in the signal. The highest frequency component in this case is equal to the maximum HR of the rodent. The maximum HR for mice typically falls within the range of 800 to 1000 beats per minute (bpm), whereas for rats, it is around 600 bpm. Based on these factors, a frame rate of 40 frames per second (fps) is suitable for contactless HR measurement in rodents, as it allows HRs to be measured up to 1200 bpm.

#### Video Compression

Assessing HR from recorded video data presents a persistent challenge, particularly for applications that are highly susceptible to external interference. In order to obtain accurate results, it is essential to have a plethysmographic signal with a sufficiently high initial signal-to-noise ratio (SNR). However, modern video encoding techniques, such as H.265, have been found to have a negative impact on the SNR, making the use of raw camera recordings necessary. This is especially crucial for rodents, where the plethysmographic signal is inherently weak. It should be noted that video compression can lead to loss of information and image quality degradation by discarding certain parts of the video data to reduce file size. This can result in the loss of important details needed for HR extraction, further emphasizing the importance of using uncompressed or minimally compressed videos for accurate HR assessment.

### Requirements for respiratory rate assessment

#### Camera positioning

While it is possible to detect movements of the thoracic and abdominal regions from a frontal perspective, there is a greater likelihood of obstruction from other animals or bedding material. In the bird-eye top view perspective, obstructions are less likely, but the thoracic and abdominal regions are not directly visible. Instead, the outer areas of the rodent body, which also move in correlation with respiration, can be used. Therefore, it is recommended to use both perspectives for respiratory detection.

#### Camera sensor

Typically, detecting movements does not require specialized camera sensors as long as the relevant areas are generally visible. Therefore, the same NIR cameras with external illumination used for HR assessment can also be used for movement detection. The use of thermal cameras is also possible and poses the additional advantage that optical tracking of rodents in the thermal range is significantly easier and more robust than in the NIR range. However, the thermal camera must have a sufficiently high resolution to detect movements at the appropriate distance.

#### Frame rates

The sampling rate theorem, discussed for HR assessment, also applies to respiratory rate measurement. The main difference lies in the frequency range, where maximum RR are not expected to exceed 400-600 breaths per minute. Therefore, frame rates of about 20 fps are sufficient for RR measurement, corresponding to a maximum detectable frequency of precisely 600 breaths per minute.

#### Video Compression

Modern video compression methods are designed according to the functioning of the human eye and visual cortex. As a result, the compression preserves motion and form better than differences in color channel intensity. This allows for detecting movements caused by respiration, even in moderately compressed videos.

### Requirements for body and core temperature assessment

#### Camera positioning

For the assessment of surface temperature, it is necessary to have the largest possible area of the rodent visible. For this, the bird’s-eye perspective is most suitable. The frontal perspective is the most appropriate for remote core temperature measurements since it corresponds most accurately to the orbital temperature. Therefore, both positions are necessary for monitoring skin and core temperatures.

#### Camera sensor

To assess temperatures, any radiometrically calibrated thermal camera sensor capable of returning thermal values within the range of +20 to +60 degrees Celsius is suitable.

#### Camera resolution

The resolution mainly determines a thermal camera’s price. While higher resolution might lead to more precise measurements, the increased price may impede the transition from the research phase to an actual product.

To estimate the animals core temperature, the eyes must be distinguishable in the thermal image. C57BL/6 and BALB/c mice have a usual body length of around 7 cm 31. The size of their eyes is approximately 2.5 mm, and roughly four pixels per eye (2 pixels on each axis) are needed to distinguish them from other facial structures. Hence a camera resolution of around 1 px/mm is needed.

### Requirements for tracking movement and circadian rhythm

#### Camera positioning

To ensure continuous tracking of rodents, they should always be visible to the camera. Therefore, the camera must be positioned to capture the entire cage area. Once again, the bird’s-eye perspective is here the most suitable option.

#### Camera sensor

The principles proposed for respiratory movement tracking also apply to general movement tracking, thus necessitating either NIR or thermal cameras.

### Housing Cage

Using standard cage bases like the EUROSTANDARD III H for mice and IV for rats offers several advantages. The cage base is the only part of the system that comes into direct contact with the animal, making using standard hardware a logical choice that has been proven over decades. These cage bases are autoclavable, rack-mountable, robust, and cost-effective. However, the cages have two main downsides: their walls are not transmissive for LWIR, and food and water are typically through a metallic mesh above the animals. In order to solve these problems, slight modifications need to be made to the cages. One is to cut a small window into the cage’s wall, which allows for capturing frontal thermal images of the animals. However, care must be taken to protect this weak point from being permanently nibbled on by rodents. Additionally, food and water should be provided in a manner that does not interfere with the functioning of top-down imagers.

### Video Acquisition and Data Storage

Computers with appropriate sensor interfaces, such as USB, CSI, or GigE, are required for controlling the cameras and acquiring video data. The hardware must be powerful enough to handle the large amount of data generated by the raw data recordings at the required frame rate, while also being compact to save space. Additionally, the hardware must be capable of real-time movement analysis to obtain accurate vital sign measurements from resting rodents. A storage system with sufficient write speed is also essential for saving the video data. Timestamps of the acquired frames must be captured alongside the image data to enable comparison with other monitoring systems. Finally, the hardware system must function as a user interface to enable animal caretakers to efficiently control the entire system.

### Handling

Ensuring the safety and welfare of animals and the accuracy and reliability of recorded data requires careful consideration of handling and disinfecting the system components. It is essential that all components that come into contact with rodents can be easily removed and cleaned with suitable disinfectants or by autoclaving. The removal process should be simple and easy to perform to ensure acceptance by animal caretakers and not interfere with existing workflows. Additionally, the system should remain robust to prevent the cleaning process from affecting the recordings.

### Homce cage system

#### Camera selection and integration

For the proposed home cage, top-down and frontal camera perspectives were used. A FLIR BOSON 320 50 FOV thermal imager (Teledyne FLIR LLC, Wilsonville, Oregon, United States) was selected for the top-down perspective. A resolution of 0.5 px/mm and complete coverage of 2000P cage floor is achieved from a 59 cm distance. The camera was used for tracking skin temperature and respiratory assessment (based on thorax expansion and contraction during the respiratory cycle). In addition, a Pi Camera Module 2 noIR, also known as PiCam 2 noIR (near-sensitive RGB camera from the Raspberry Pi Foundation, Cambridge, England, UK), was used for activity monitoring (including tracking) and cardiac assessment. Hence, enclosures for both cameras were designed to be permanently attached to them. The enclosures were manufactured on a regular 3D printer. Fig 2 displays the Raspberry Pi Camera and FLIR boson enclosures created for this project.

**Fig 2.**
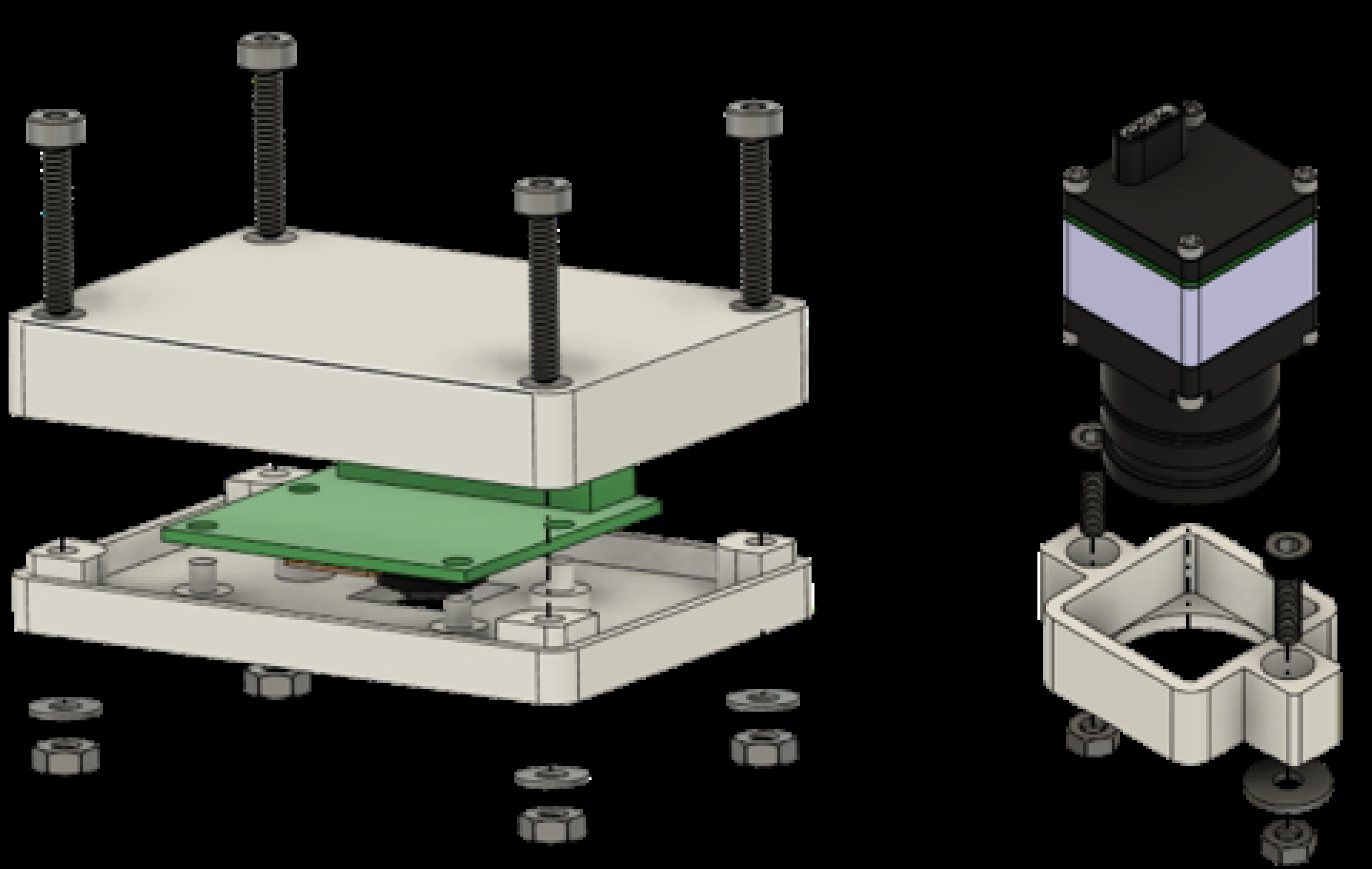
Camera assemblies at the top of the cage. Displayed on the left is a PiCam 2 noIR module in its enclosure. The right assembly shows the mount for an FLIR Boson 320 with 50 FOW

A FLIR Lepton 3 (a lower-resolution thermal imager from Teledyne FLIR LLC, Wilsonville, Oregon, United States) was chosen for the frontal perspective. Its minimal focus distance of 15 cm achieves a resolution of 0.75 px/mm, which is just below the requirement. Unfortunately, this is the only cost-effective thermal imager with a shorter image distance. This camera was applied for core temperature estimation and respiratory assessment. An additional Pi Camera Module 2 noIR (Raspberry Pi Foundation, Cambridge, England, UK) is used for grimace scale and cardiac analysis. The cameras are next to the water bottle to increase the chance of good shots of approaching rodents. Because polysulfone (the material of the cage base) is opaque to thermal radiation, a window needs to be cut into it. Indeed, this might introduce a weak point for nibbling. To avoid that, the edges of the window were protected with a metal eyelet/grommet (with an inner diameter of 16 mm). To protect the thermal cameras lens from liquids and teeth, an opaque 0.1 mm thick IR transmissive film (Kube Electronics AG, Gossau, Switzerland) was placed before it. The color camera was slightly tilted in its mount to increase the FOV overlap of both imagers (see Figure 3). Because the cage base needs to be autoclavable, the cameras were not permanently fixed to the base. Instead, M3 t-slot nuts were installed at the base to easily attach the cameras by sliding them in.

**Fig 3.**
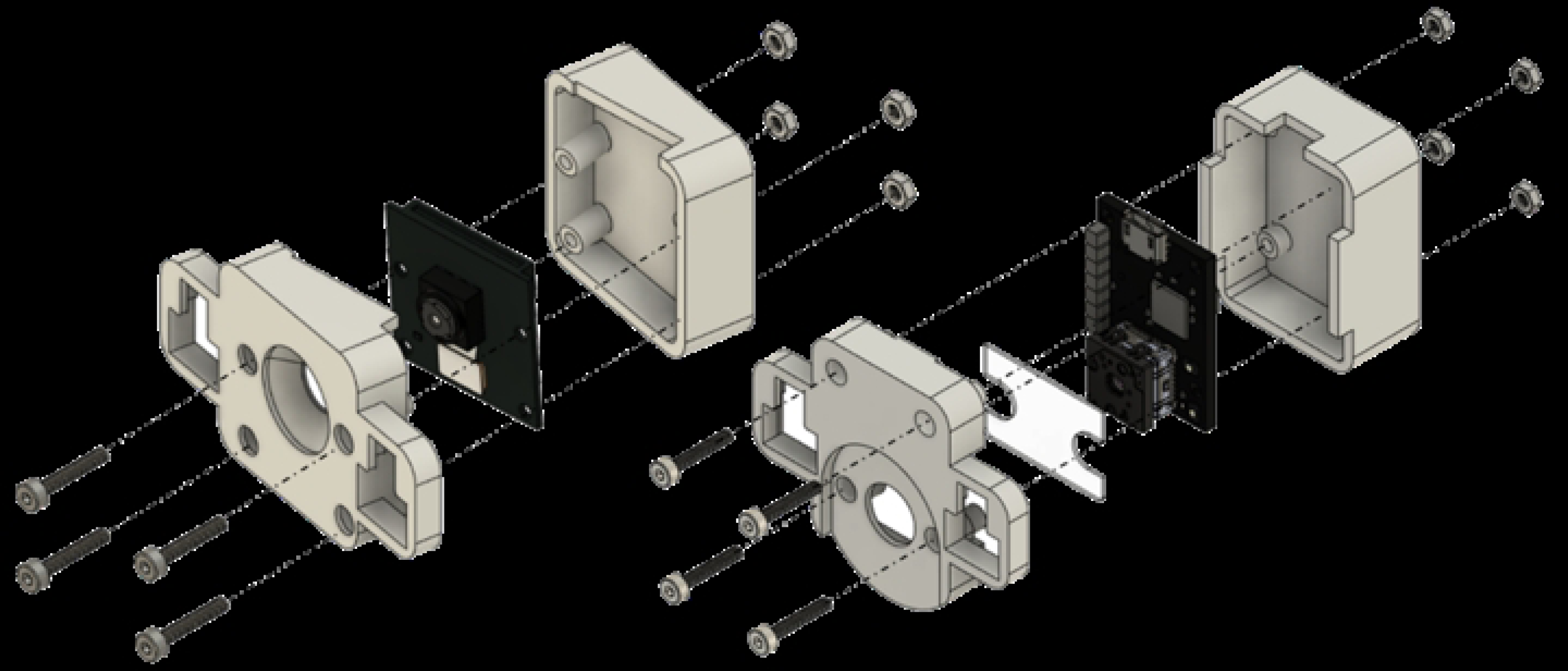
Frontal camera assembly near the water source. The left assembly contains a PiCam 2 noIR module in its enclosure. The right assembly holds a FLIR Lepton 3 camera module for thermal images.

Table 1 provides a comprehensive overview of the cameras employed in the home cage system. It highlights the purpose of these cameras, the resolutions at which they operate, and their specific installation positions.

**Table 1.** Cameras of the monitoring cage.

| Camera | Purpose | Resolution (px) | Location |
| --- | --- | --- | --- |
| FLIR BOSON 320 50° FOV | thermal imager for tracking and respiratory assessment | 320x240 | Top-down view of the whole cage |
| Pi Camera Module 2 noIR | near infrared sensitive RGB camera for activity and cardiac assessment | 1920x1080 |  |
| FLIR Lepton 3.5 | thermal imager for core temperature estimation and respiratory assessment | 160x120 | Frontal view next to the water bottle |
| Pi Camera Module 2 noIR | near infrared sensitive RGB camera for grimace scale and cardiac analysis | 1920x1080 |  |

### Cage design

The cage design was driven by the required cage bases (EUROSTANDARD III H for mice and IV 2000P for rats, respectively) and the optical parameters of the imagers. The height of the cage was calculated so that the FOV of the thermal imager and Pi Camera exactly covered the complete cage floor. The width and depth were directly derived from base dimensions (612×435×216mm). Two designs were created: one for a cage that can be rack mounted and one for a standalone cage, which can also be used with telemetry plates (Fig 4).

**Fig 4.**
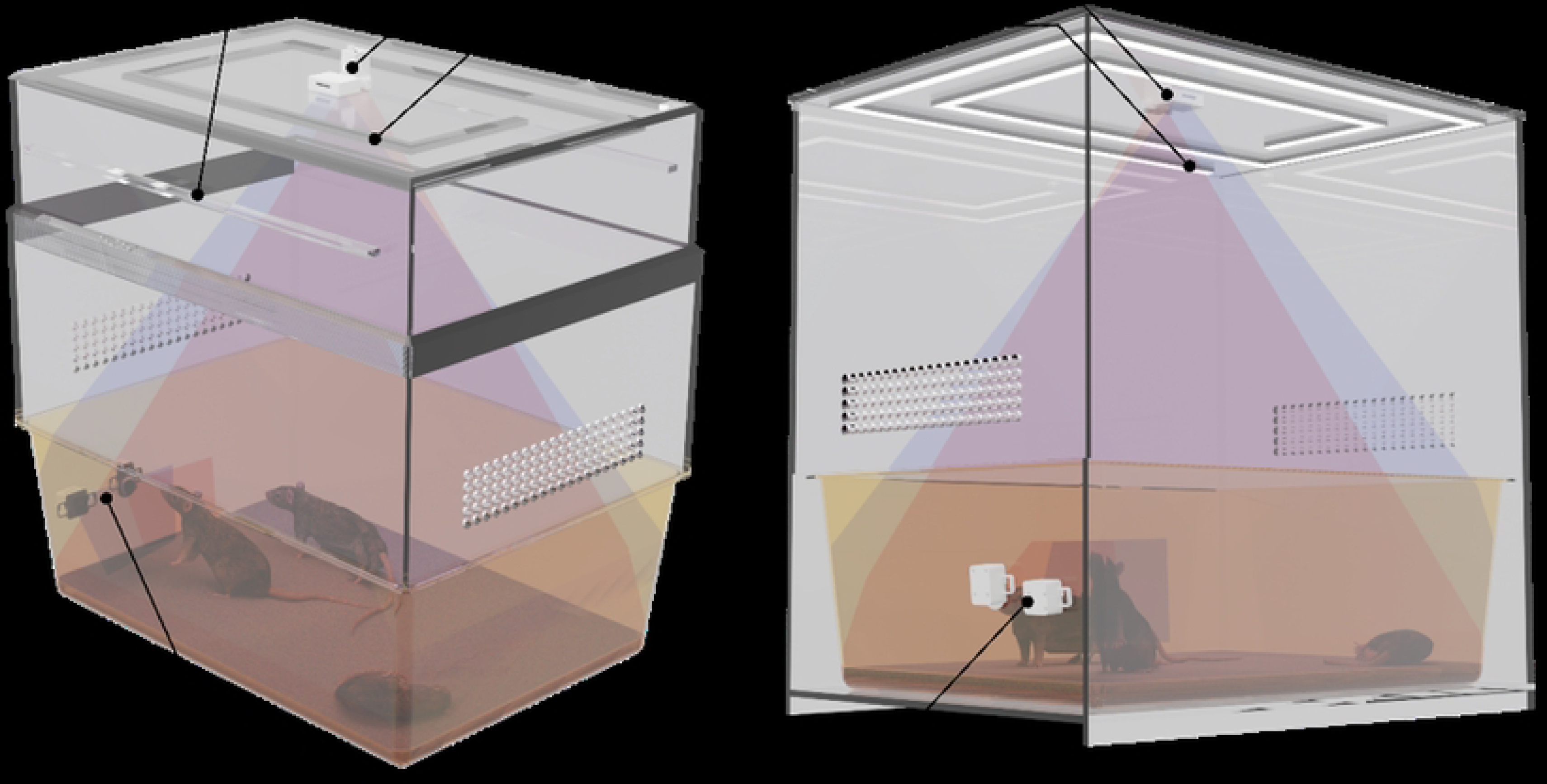
Home cage designs. The left design is intended to be used in a cage rack, while the right design is for stand-alone settings.

In both cases, the cage bases can easily be pulled out after removing the frontal cameras to allow for animal handling. While for the rack mount approach, the top of the cage remains inside the rack supported by dedicated rails, the standalone cage features a “drawer slide” where the base can be pushed into the outer enclosure. To ensure adequate lighting 850nm NIR led strips have been mounted to the cage top in spirally arranged LED profiles with diffusers. The cage enclosure is made from transparent acrylic sheets with a thickness of at least 7 mm. The 3D models of the two prototypes are provided as supplementary materials.

### Monitoring hardware implementation

To satisfy the system requirements, a customized software and hardware system was developed to facilitate the connection of the cameras and the management of the data recording. A detailed illustration of the hardware concept is presented in Fig 5.

**Fig 5.**
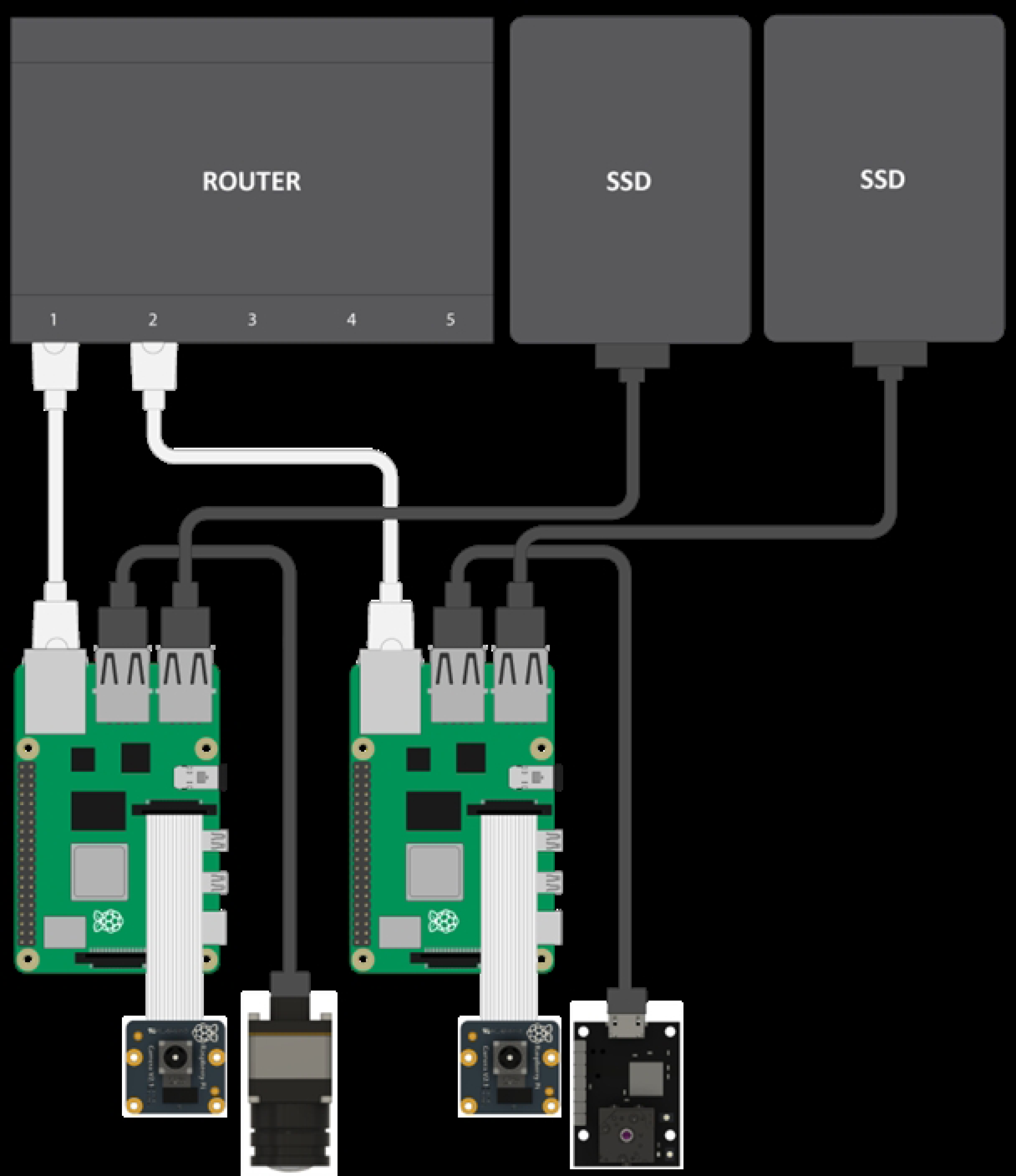
Illustrated hardware concept. The hardware consists of a dedicated network router (a), fast SSD data storage (b), two Raspberry Pi 4B computing modules (c) each connected to a Raspberry Pi Camera Module v2 NoIR (d), and a Flir Boson 320 (e) and Flir Lepton v3.5 (f) thermal camera respectively.

Raspberry Pi 4 B single-board computers (Fig 5c) have been selected as the recording hardware for our concept implementation.The selected Pi cameras (Fig 5d) can be controlled natively via the integrated CSI interface and the FLIR cameras (Fig 5e-f) can be controlled in parallel via USB 3.0. Additionally, the acquired data can be stored on external hard drives via USB (Fig 5b) or on a network-attached storage (NAS) via Gigabit Ethernet. Furthermore, these devices are reasonably priced and have a wide range of software and programming language support.

In our setup, a system was implemented that utilizes two separate modules, one dedicated to the cameras located in the lid and the other for the cameras in the frontal area of the enclosure. This separation ensures an even distribution of the system load and acquired data between the modules, which is essential for processing and transfer, given the large amount of data generated by the high frame rate recordings. However, even with modern recording hardware, this presents a significant challenge, as the theoretical performance of CSI or USB is often limited by the processing interface, making achieving the desired rates a non-trivial task. Therefore, a customized software solution based on Python and C++ was developed for system management and data recording for the modules. This approach allows low-level access to the cameras and can use their maximum FPS capabilities, i.e., 40 FPS for the Picamera v2 and 60 FPS for the Boson. The implemented system utilizes two separate storage modules, one for the cameras in the lid and the other for the frontal cameras. An even distribution of the system load and acquired data between the modules is ensured by this separation, which is essential for processing and transfer, given the large amount of data generated by the high frame rate recordings. In addition, a recording protocol based on Google’s Protocol Buffers (Protobuf) was developed to enable the synchronization and the integration of further monitoring modules or external sensors.

### User interface

Its user interface is essential to consider in implementing the measurement system. A user-friendly interface prevents errors resulting from misoperation and increases user acceptance. In addition, it is crucial that the system supports all standard operating systems and can be integrated into existing infrastructures. To address these requirements, a web application-based solution (Fig 6) was developed, allowing for remote control of the recording process. This solution is facilitated through an integrated web server, accessed through the Raspberry Pi’s built-in WiFi access point or a cable-based network connection. This approach allows easy access to the system from any device with a web browser, including smartphones, tablets, laptops, and desktop computers.

**Fig 6.**
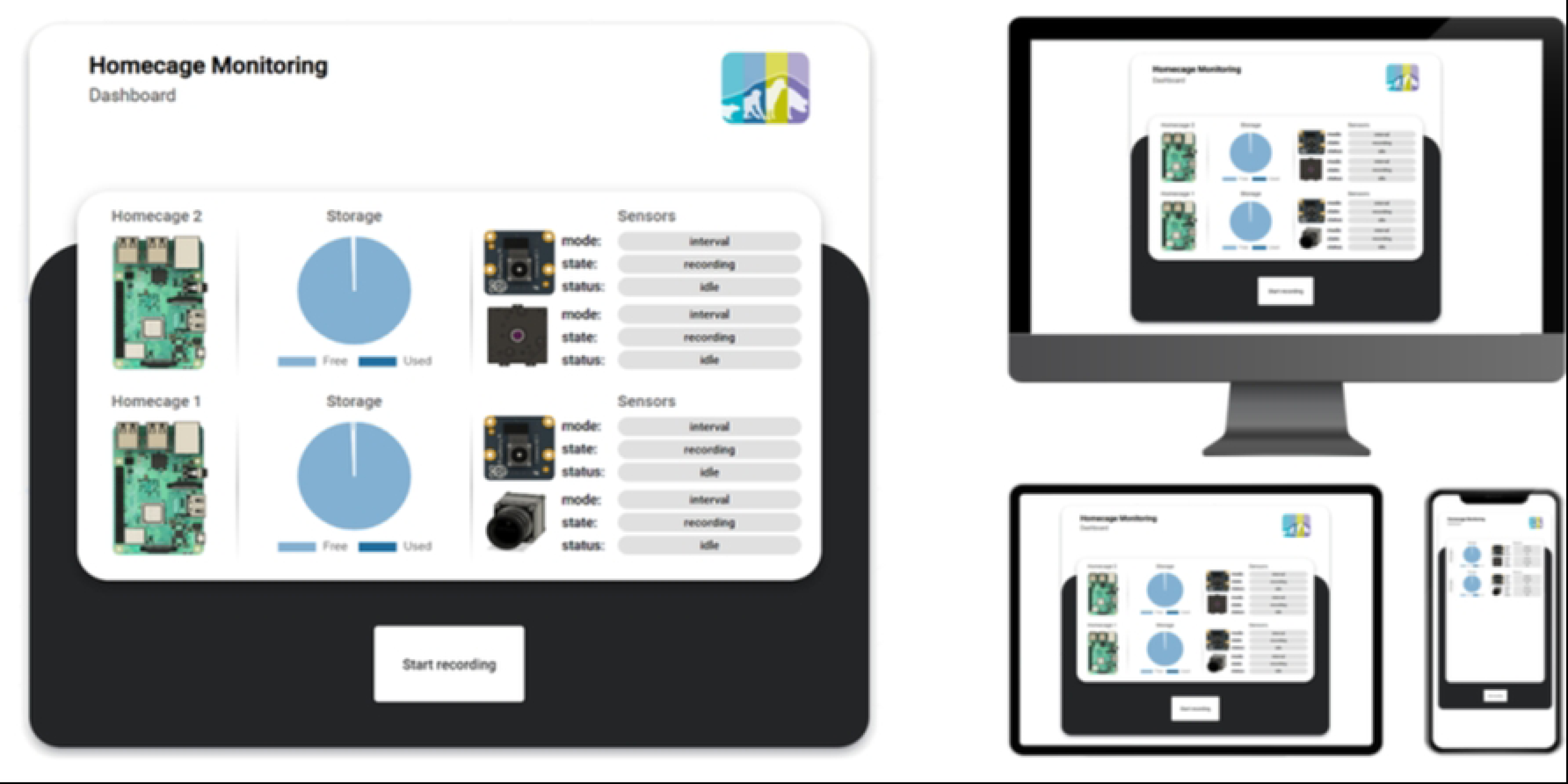
User Interface. One-Button design concept, providing an overview of the systems and system sensors status (left) and enabling multi-device access (right).

### Resources

The paper provides open access to all the specifications, software, system configurations, and 3D models presented. Researchers and interested individuals are encouraged to access and utilize these resources for their own studies and experiments. By visiting the GitHub repository at the following link, users can explore and download the provided materials: https://github.com/FOR2591/homecage

### Experimental evaluation

#### Animals

In this study, four adult, male C57BL/6 mice were used. The mice were housed inside our proposed system with a euro standard type III cage as the base, with sawdust bedding, nestlets, and a tube. The room temperature was maintained at 22 *±* 2*^◦^* C, with a relative humidity of 40-60 and a 12-hour light-dark cycle.

#### Ethical statement

The animal experiment conducted adhered to the guidelines set by the German federal law concerning animal protection ad use for scientific purposes. The animals used were part of an setting for further education and advanced training and received authorization from the animal welfare committee of the government (LANUV, North Rhine-Westphalia, Germany, AZ: 81-02.05.40.18.0537). Animal housing and care complied with both institutional regulations and EU Directive 2010/63.

#### Study Design

Feasability tests were conducted at the Institute for Laboratory Animal Science and Experimental Surgery, Faculty of Medicine, RWTH Aachen University, Aachen, Germany, to verify the overall functionality of the proposed system, including the chosen cameras, lighting, and camera positioning. Prior to their regular interventions, the test group of four mice was transferred together to the monitoring system.

Recording started approximately one hour later, allowing them to acclimate to their new environment. The mice were then monitored for two weeks. Recordings were conducted with and without artificial daylight to assess the impact of lighting conditions on the measurements.

## Results

To validate the quality of the obtained recordings, as a first step, a visual inspection was performed to approve the pre-calculated angles and mounting points of the cameras concerning the mice, as illustrated in Fig. 7. Concerning the cameras mounted on the lid, it can be seen that the pre-calculated height is sufficient to cover the entire bottom area as well as the walls of the inner cage tray, allowing the animals to remain visible even when climbing up the edges, for example, on the feed trough or the enrichment. In the frontal recordings, we observed that the camera’s field of view is optimally utilized as long as the bedding is at a normal level and the mice are near the camera. However, when the mice begin to dig in the bedding, a large part of the image may be obscured by the bedding (see fig. 7 thermal front). Therefore, it is necessary to consider adjusting the height of the frontal cameras depending on the extent of the digging behavior.

**Fig 7.**
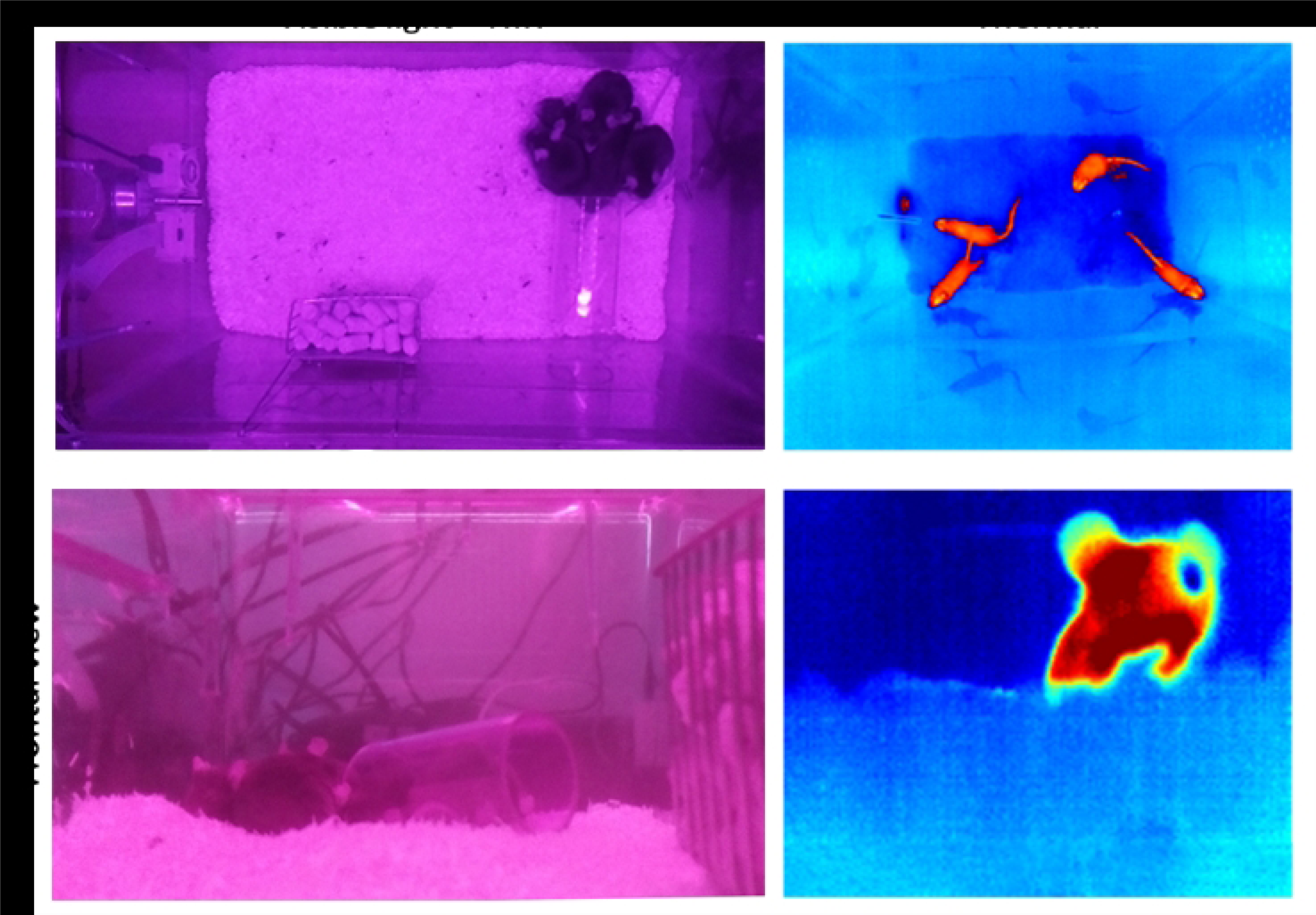
Example images of cameras. Captured images of all four integrated cameras, which are segregated into top and frontal views as well as visible light and NIR and thermal views.

To evaluate the upper Pi camera’s ability to monitor vital parameters, a sequence of 15 seconds was selected from the recording with active daylight illumination and without active daylight illumination, during which all animals remained motionless, thereby eliminating motion artifacts as a source of error. Within these 15-second recordings, three separate ROIs were defined, and their respective signal spectra were analyzed. The position of these ROIs is shown in Fig. 8 and Fig. 9, and they included an unfurrowed skin area located in the ear of a mouse (1), an area along the outer edge of the mouse (2), and a reference area that did not contain any mice (3).

**Fig 8.**
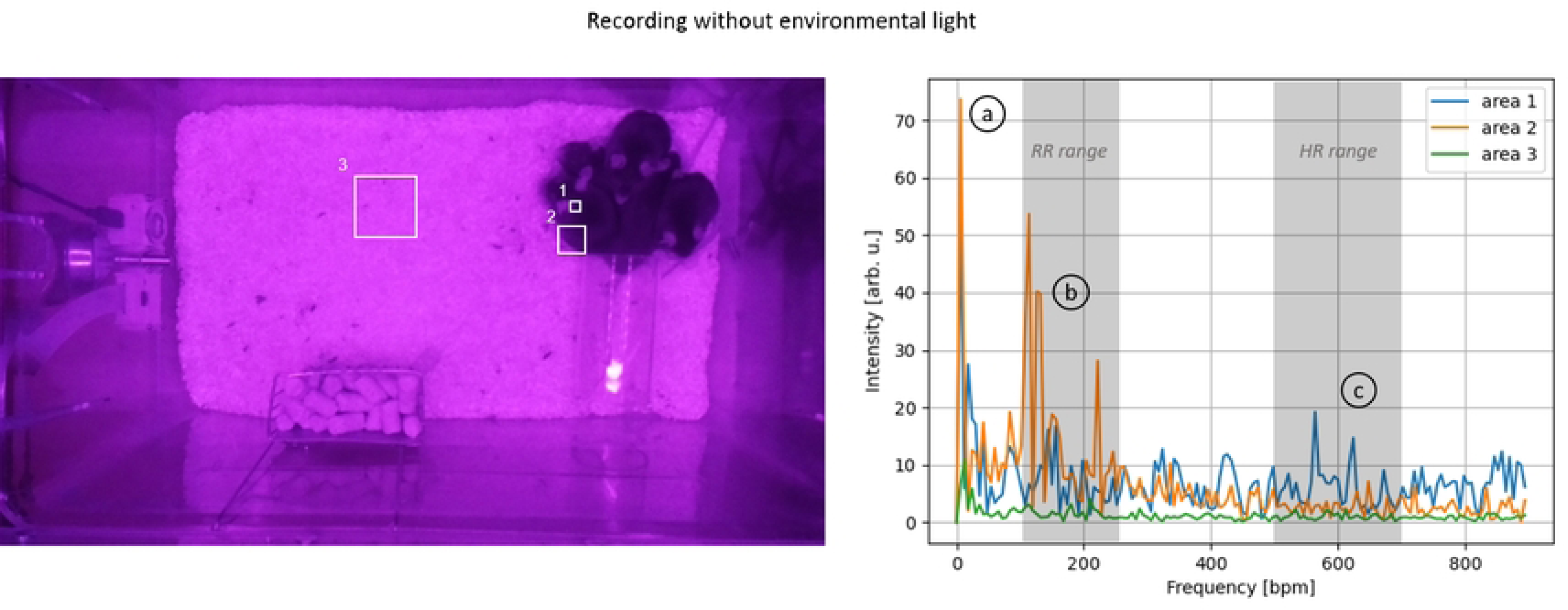
Spectral investigation without environmental lighting (top) (1) Inner ear area, (2) Mice’s outer edge, and (3) Mice-free area. The spectral analysis is shown on the right side, and the left side displays the corresponding areas. The spectra indicate notable harmonics at three different ranges: (a) 6 bpm, (b) 100-250 bpm, and (c) 400-600 bpm.

**Fig 9.**
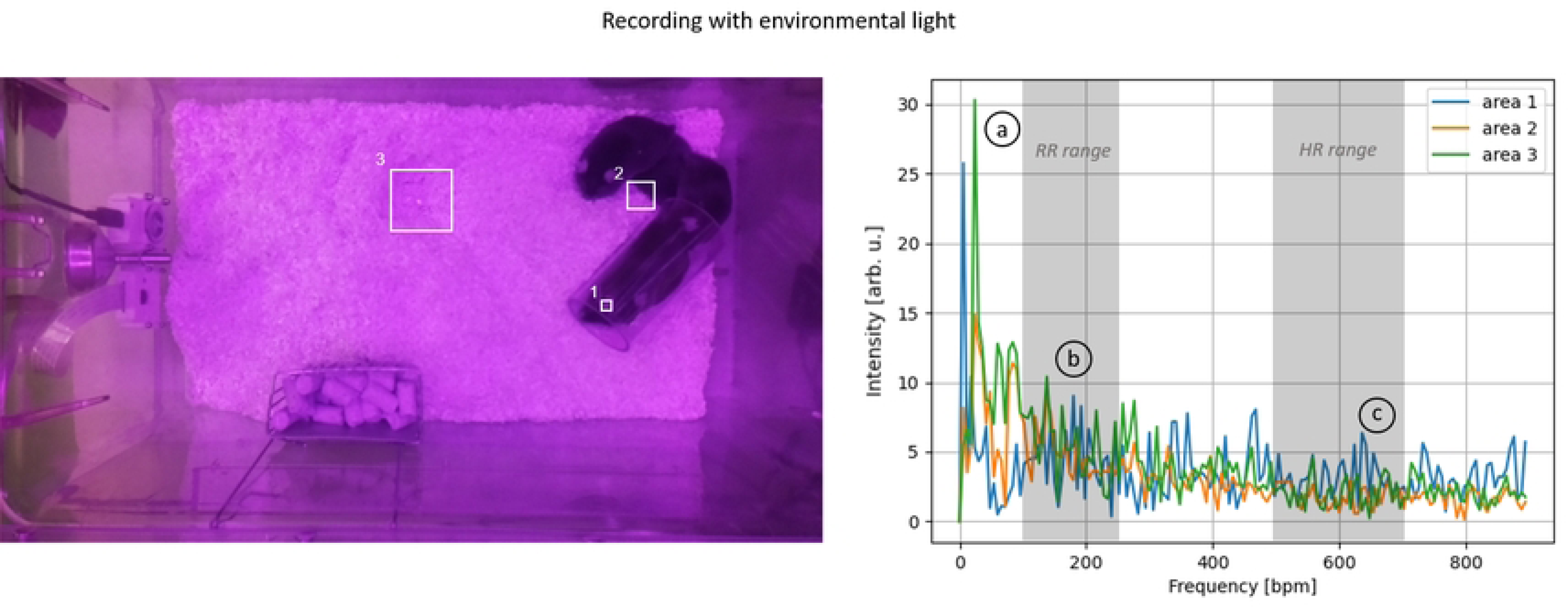
Spectral investigation with environmental lighting (top) (1) Inner ear area, (2) Mice’s outer edge, and (3) Mice-free area. The spectral analysis is shown on the right side, and the left side displays the corresponding areas. The spectra indicate notable harmonics at three different ranges: (a) 6 bpm, (b) 100-250 bpm, and (c) 400-600 bpm.

Without the illumination of the room light (Fig. 8), the signal intensity changes in the reference region (area 3) were significantly smaller than those caused by the mice (areas 1+2). Thus, no adverse effects on measuring vital parameters are to be anticipated. All areas showed a prominent first harmonic at approximately six bpm. This prominent harmonic was less pronounced in the reference area than in the other two. This observation suggests that this harmonic may not be attributed to the influence of illumination but rather to a general oscillation in the room, especially when compared to the recording with illumination where this harmonic is also present. The region along the edge of a mouse (area 2) showed distinct first and second-harmonic oscillations in the range of 100 to 200 breaths/min, corresponding to the typical range of respiratory activity. Inside the ear (area 1), where changes in blood flow can be observed, a distinct first harmonic could be identified, corresponding to the typical HR range of 400-600 bpm.

Under ambient lighting conditions (Fig. 9), the obtained recordings showed similar spectra to those without ambient light. In both cases, the first harmonics were distinct in the ranges corresponding to the HR and RR. However, the spectra of all three areas demonstrated a markedly lower ratio between the harmonics and the noise floor, thus to a worse SNR. This observation indicates that ambient lighting has an adverse effect on the measurement of vital parameters.

Analogous investigations were performed with the frontal Pi camera. Again, 15-second recordings were analyzed with and without environmental lighting, focusing on areas 1-3, corresponding to the regions captured by the upper camera but from a frontal perspective. In this case, we can observe similar peaks for RR and HR in the relevant ranges (Fig 10, b-c). However, the frontal recordings exhibit a significantly higher noise level within and between the areas, resulting in poorer SNR overall. This applies to both RR and HR. Additionally, from the frontal perspective, the low-frequency noise component of the system is noticeable aswell.

**Fig 10.**
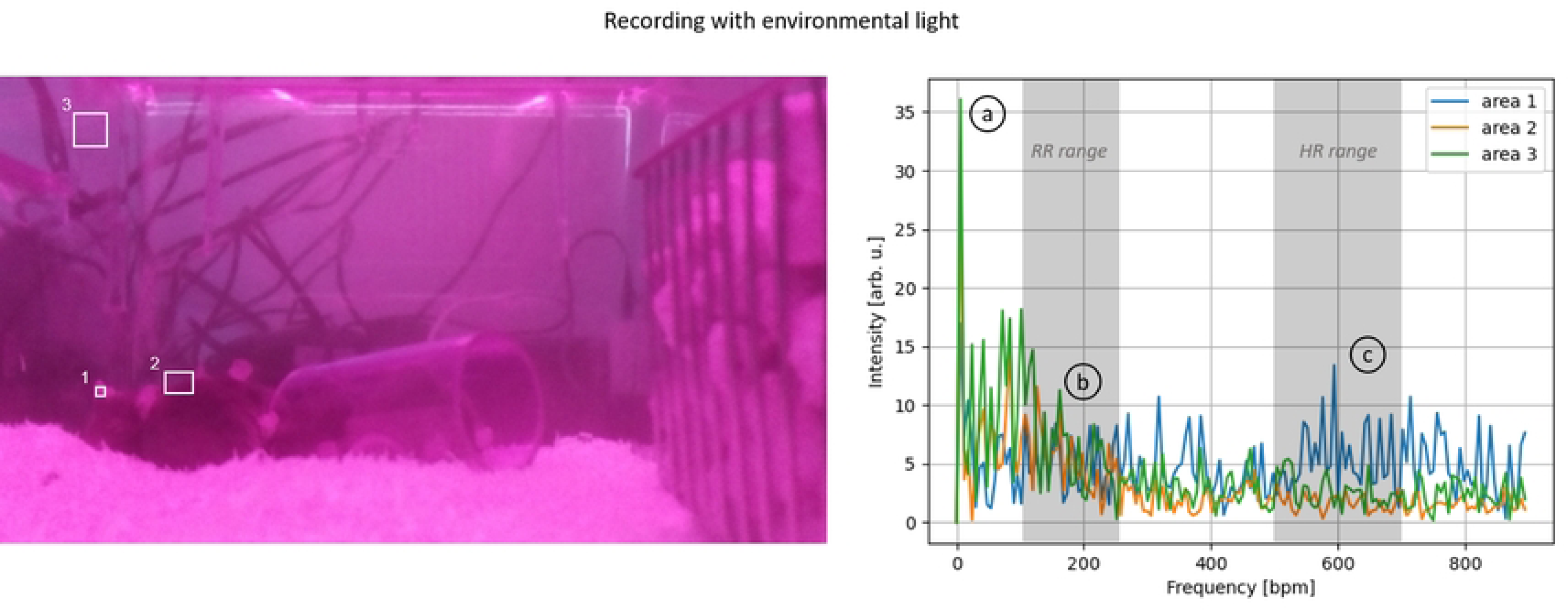
Spectral investigation without environmental lighting (front) (1) Inner ear area, (2) Mice’s outer edge, and (3) Mice-free area. The spectral analysis is shown on the right side, and the left side displays the corresponding areas. The spectra indicate notable harmonics at three different ranges: (a) 6 bpm, (b) 100-250 bpm, and (c) 400-600 bpm.

The same evaluation with room light reveals a similar pattern (Fig. 11). However, the spectra of the individual areas are more noisy in contrast to the frontal recordings without room light, particularly in reference area 3. Moreover, the respiratory pattern is less identifiable in the spectrum due to lesser contrast.

**Fig 11.**
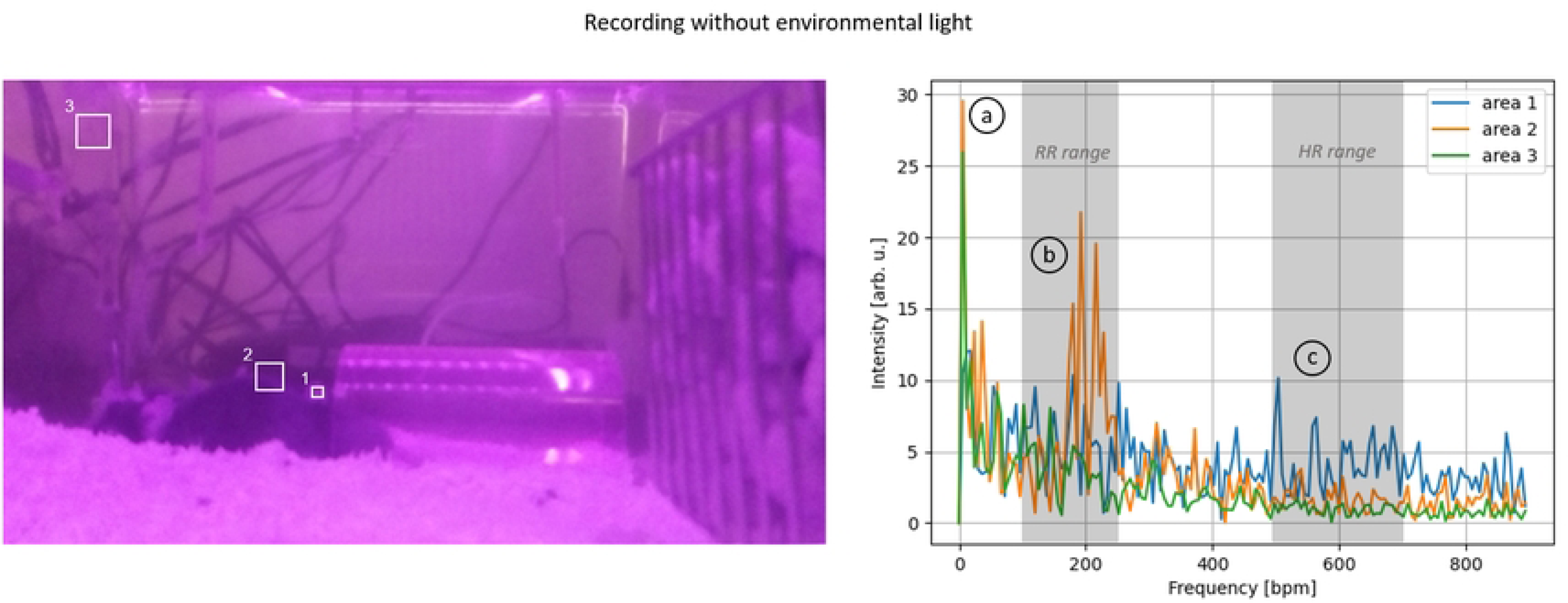
Spectral investigation with environmental lighting (front) (1) Inner ear area, (2) Mice’s outer edge, and (3) Mice-free area. The spectral analysis is shown on the right side, and the left side displays the corresponding areas. The spectra indicate notable harmonics at three different ranges: (a) 6 bpm, (b) 100-250 bpm, and (c) 400-600 bpm.

## Discussion

This paper proposed a concept of a smart home cage monitoring system for mice, which is also adaptable for rats. It is the first study that addresses the available algorithmic approaches, technical requirements, and implementation. The concept and the prototypes were developed during the “FOR 2591: Severity assessment in animal-based research” project funded by the German Research Foundation.

Although the primary focus of this project relies on the unobtrusive assessment of vital parameters, the great advantage of this concept is that it can also be used for monitoring 24/7 other relevant aspects in animal experiments or breeding facilities. Furthermore, as the cameras cover the entire cage, physiological and non-physiological parameters can be assessed simultaneously and continuously using the same sensors, which has proven relevant for animal-based research [35, 36].

The smart home cage system presents a modular design that allows researchers to add cameras according to their needs and the parameters they want to monitor. This modularity makes the system adaptable to different experimental designs and provides flexibility for future upgrades. Additionally, the positioning of the cameras provides a complete top view of the cage, enabling monitoring of the animals’ activities, vitals, and circadian rhythm, as well as a close view of the animals around the drinking bottle.

Significant peaks in the measured spectra were identified during the initial trials on mice using the system, suggesting that the estimation of HR and RR, which are challenging parameters to obtain, is feasible. Similar results are expected for rats, as their larger size and slower HR and RR make the extraction process inherently easier. Future publications will provide a complete proof of concept regarding vital parameters, motion, and circadian rhythm.

Current monitoring modalities, more properly telemetry systems, have several disadvantages. First and foremost, the implantation can be stressful and painful for the animal, potentially causing discomfort or injury [37]. Additionally, the implanted device may interfere with the animal’s natural behavior, movement, and ability to interact with its environment. Moreover, it can have considerable unfavorable physiological impacts. Thus, replacing these methods with unobtrusive techniques fulfills the refinement principle as proposed by Russell and Burch. Additionally, the number of animals that can be monitored with a camera-based system is not inherently limited, as any number of animals can be monitored simultaneously. This represents a significant advantage over systems from well-known telemetry manufacturers, where simultaneous monitoring multiple animals is impossible, complex, or associated with considerable additional costs. Moreover, group housing should be the default housing method since rodents are inherently social animals. Keeping them in isolation has proven to negatively affect their physiology and behavior [38–40], which stands against the 3R guidelines for animal welfare and impedes the validity of experimental results [41]. Furthermore, using a transmitter may increase the risk of infection or other complications, which can be avoided by employing contactless methods that also enable repeated measurements without compromising animal welfare and do not require batteries.

Stressful conditions during animal studies can negatively impact their health, behavior, and experimental outcomes, with handling and restraint being a common sources of stress. Inappropriate and harmful handling can cause fear, anxiety, and physical harm [42]. By using a monitoring home cage, such harmful handling can be significantly reduced.

Several other advantages come along with such revolutionary technologies. For example, contactless measurement of vital signs allows for continuous monitoring of animals’ vital signs over extended periods without disrupting them or the experimental conditions, providing more accurate data and reducing the number of animals required for experiments. Moreover, objective parameters such as HR, respiratory rate and rhythm, temperature, and circadian rhythm alterations, among other behavioral parameters, can be used to create a score similar to the Early Warning Score (EWS) used in human medicine or to integrated into existing scores like RELSA [13]. Such scores can help researchers assess and take early measures if the animal’s health deteriorates and revolutionize animal research by providing a deeper understanding of disease progression.

Such digitized monitoring cage systems also offer the possibility of a centralized overview of multiple cages at once. In addition, researchers can collect data on all individual cages within a rack remotely. Cameras also improve staff safety by enabling remote animal behavior monitoring, particularly when animals pose a risk. For example, when working with models of infectious disorders.

Although a monitoring cage has enormous potential, some drawbacks must be discussed. First, as it is almost impossible to differentiate between the animals due to their almost identical appearance, it is necessary to establish a method of differentiation, such as marking the tail of each animal with rings. Secondly, continuous assessment of vital parameters may be impeded by movement artifacts or if animals are obscured by nesting material, being burrowed, or hidden inside enrichment such as tubes. Thirdly, the current 3D models were developed primarily for the 1290D EUROSTANDARD TYPE III cage from TECNIPLAST S.p.A. (Buguggiate, Italy). Hence, adjustments are necessary to apply this method to different cage models. These adjustments include recalculating the cage height to ensure complete coverage of the cage bottom within the camera’s field of view. Fourthly, a central data storage system must be developed to archive data from all rack cages. However, the main challenge here lies in handling the sheer amount of data; therefore, preprocessing methods must be implemented to reduce the data before archiving. In addition, it must be considered where the data should be analyzed, either centrally in one high-level performance computer or individually (i.e., one computer for each cage). Finally, even though there have been significant developments in assessing vital parameters using cameras, they are incapable of measuring EEG or ECG. Therefore, for experiments studying cerebral activity or heart diseases, the implantation of telemetry sensors may still be necessary.

## Conclusion

The use of monitoring cages and cameras for assessing animal welfare based on non-physiological parameters such as food intake, behavior, social interaction, and movement has shown great potential in recent years. However, to date, no remote monitoring system has been capable of measuring physiological parameters. This paper presented a new concept for a monitoring home cage, capable of monitoring physiological and non-physiological parameters through unobtrusive monitoring. Such unobtrusive monitoring of an animal’s behavior and vital parameters can significantly impact animal-based experiments by enabling researchers and animal caretakers to detect and respond to changes that may indicate poor welfare. As technology continues to advance, these methods are likely to become even more sophisticated and accessible, contributing to ensuring that laboratory animals receive optimal care and refinement as well as improving the overall quality of animal-based experiments. Although not all parameters can be measured with current technologies, using these methods when relevant physiological signals, such as ECG or EEG, are strictly required for the experiment or animal welfare may reduce the need for implantable sensors.

